# Effect of Lead Body and Helix Design Variables on Implantation Success, Insertion Depth, and Muscle Torque in Left Bundle Branch Area: Insights from An Ex-Vivo Porcine Model

**DOI:** 10.1101/2025.11.25.690583

**Authors:** Ankur R. Shah, Alex Puccio, Kyoichiro Yazaki, Emmanuel Offei, Martha Sofia Ruiz Castilo, Surachat Jaroonpipatkul, Ava Yaktaeian Vaziri, Muhammad S. Khan, Ravi Ranjan, Robert Hitchcock, Derek J. Dosdall

**Affiliations:** Nora Eccles Harrison Cardiovascular Research and Training Institute; Department of Biomedical Engineering; Division of Cardiothoracic Surgery, Department of Surgery; Division of Cardiovascular Medicine, Department of Internal Medicine The University of Utah, Salt Lake City, UT 84112, USA

**Keywords:** Left Bundle Branch Area, Interventricular Septum, Pacing Leads, Lead Behavior, Helix design, Ex-vivo

## Abstract

**Background:** Lumenless and stylet-driven leads used for left bundle branch area pacing differ in design and have a significant implantation learning curve. While prior studies examined longer helices for deep septal pacing, the influence of other design variables remains unclear.

**Objective:** To evaluate how helix design and axial force affect interventricular septum insertion efficacy.

**Methods:** Rigid leads were developed using helical coils with variable outer diameter, number of turns and pitch. Porcine septa (n=16) were clamped perpendicularly for insertion using an optimized rotation-response system. Axial force simulating lumenless (30g) or stylet-driven (60g) leads was applied, and a fixed number of rotations were delivered at a constant rate. Each helix design (n=8) was tested 3x per axial force at three septal sites. Insertion depth, muscle-torque and visual feedback were recorded. Insertion was successful if depth exceeded coil length without surface entanglement. Effects of design factors were compared.

**Results:** At 30g, more helix turns significantly improved insertion success (P=0.04), while fewer turns frequently produced entangled failure (P=0.04) marked by high torque variability (P<0.001). Smaller-pitch helices trended toward higher torque and success, whereas larger pitch achieved greater depth (P=0.05). Larger outer diameters also trended toward higher torque and improved success at 30g. At 60g the influence of helix design variable diminished and consistently yielded higher than at 30g.

**Conclusion:** An optimized lead rotation-to-translation system elucidates how helix geometry and axial force interact during septal insertion. These interactions are explainable using an intuitive mechanical framework which is helpful for optimizing lead design.

## 1. Introduction

Implantable cardiac electrodes with helical coils were first introduced in the early 1960s^1^. Over the next four decades developments in transvenous pacing leads with helical screw-in electrodes addressed limitations at the electrode tissue interface to provide safe, stable, and low stimulation-threshold pacing^2^. Thinner pacing leads (4-5 Fr) developed in early 2000s have enabled physicians to attempt permanent cardiac pacing at alternate sites and treat a wider patient population^3^. Recently, permanent left bundle branch area (LBBA) pacing has emerged as a primary pacing target for bradycardia and heart failure indications^4^.

While LBBA lead implantation techniques are being standardized, the procedure is associated with a significant learning curve.^5,6^ The standard helical screw-in electrode tip has limitations in ventricular septal penetration^7^. During septal penetration, multiple lead insertion attempts are often required due to the inability of the lead to advance into the interventricular septum (IVS) or if an undesired paced morphology is observed^8^. Lead advancement may fail if the collagenous layer on the endocardium and/or chrordae tendineae wrap around the helix, leading to lead entanglement (Figure 1A). Lead advancement may also fail if myocardial tissue is too firm due to scar or extensive fibrosis to permit further advancement, creating a mechanical barrier^7^. Additionally, lead advancement may also fail if the lead tip lands in the recess between two trabeculae with no myocardium in its path^7^. This may result in a “drill effect,” where the lead can only rotate without advancing further (Figure 1B). In such situations the lead must be repositioned^7^.

**Figure 1.**
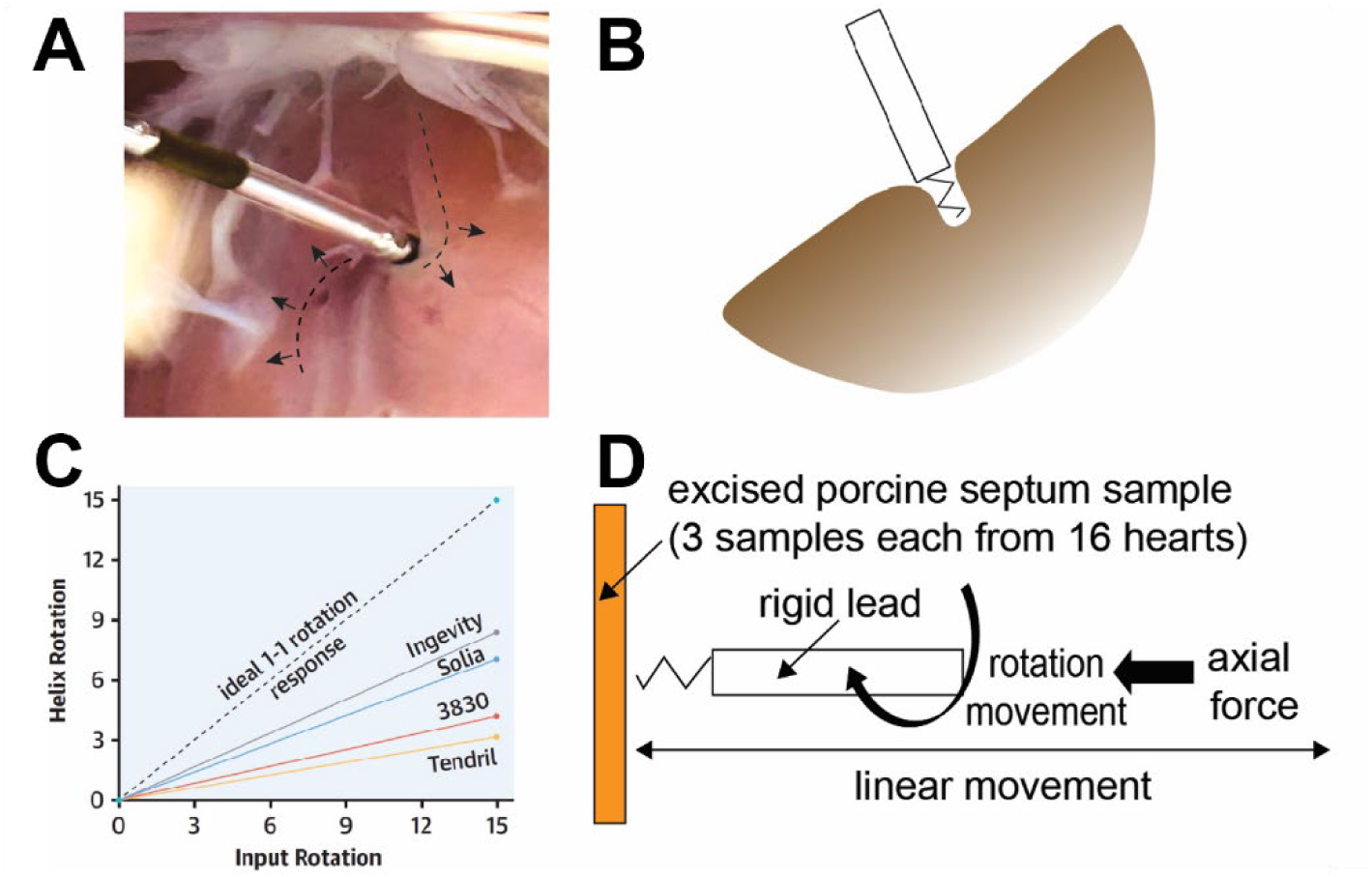
A benchtop experimental rig developed to evaluate how helix variables affect septal lead penetration in an ex-vivo porcine septum model. Common septal penetration failed modes: **(A)** lead helix ‘entangled’ in muscle (felt as torque buildup on the lead due to muscle stretching), or **(B)** helix stuck in a septal recess and unable to interact with the muscle. **(C)** In commercial leads, only a fraction of applied input rotations translates to helix rotations. This relationship is dependent on lead design. Reprinted from JACC: Clinical Electrophysiology, Vol 10, No 2, D. Chapman et al., ” Assessing Torque Transfer in Conduction System Pacing”, Pages 306-315, Copyright (2024), with permission from Elsevier. **(D)** An idealized one-one rotation response system was developed to isolate the impact of lead helix design variables while eliminating the impact of torque loss along the lead body.

Even when the lead successfully advances into the IVS, the number of rotations applied to the lead does not appear to correlate with the thickness of the septal wall due to torque buildup within the lead body^8^. Moreover, only a fraction of the rotations applied to the lead result in helix rotations and this relationship varies depending on the specific lead design (Figure 1B)^9,10^. Such variation may arise from differences in the construction of the lead body and/or the helix among the lead models used in clinical practice. While previous studies have explored longer helix coils for deep septal pacing in small cohorts^11,12^, the role of other helix design variables on IVS penetration remains to be understood.

In this study, the isolated impact of helix design variables on successful IVS implantation were evaluated. This was achieved by transmitting all the input torque to the helices using a customized rotation-to-translation motion ex-vivo setup fixed with rigid leads of different helix configurations (Figure 1C). The setup is integrated with a system to measure torque experienced by the septal muscle (muscle torque), allowing for deeper understanding of helix-tissue interactions during insertion.

## 2. Materials and Methods

### 2.1 Experiment setup

Rigid leads (RLs, development described later) with helices of variable configurations were mounted on a system that translated rotational motion to lead insertion. The system includes a pin-vise to clamp RLs (240c, Starrett, MA) with ball bearings allowing for free rotation (99502H-2RS, uxcell, CN), and a linear slider (SEBS15B, Nipponbearing, JP) connected to ball bearings to guide the linear motion. Septal tissue samples were clamped perpendicularly to the linear slider to facilitate RL insertion. A schematic and the physical setup of the idealized rotation response system is shown in **Figure 2A, 2B**.

**Figure 2.**
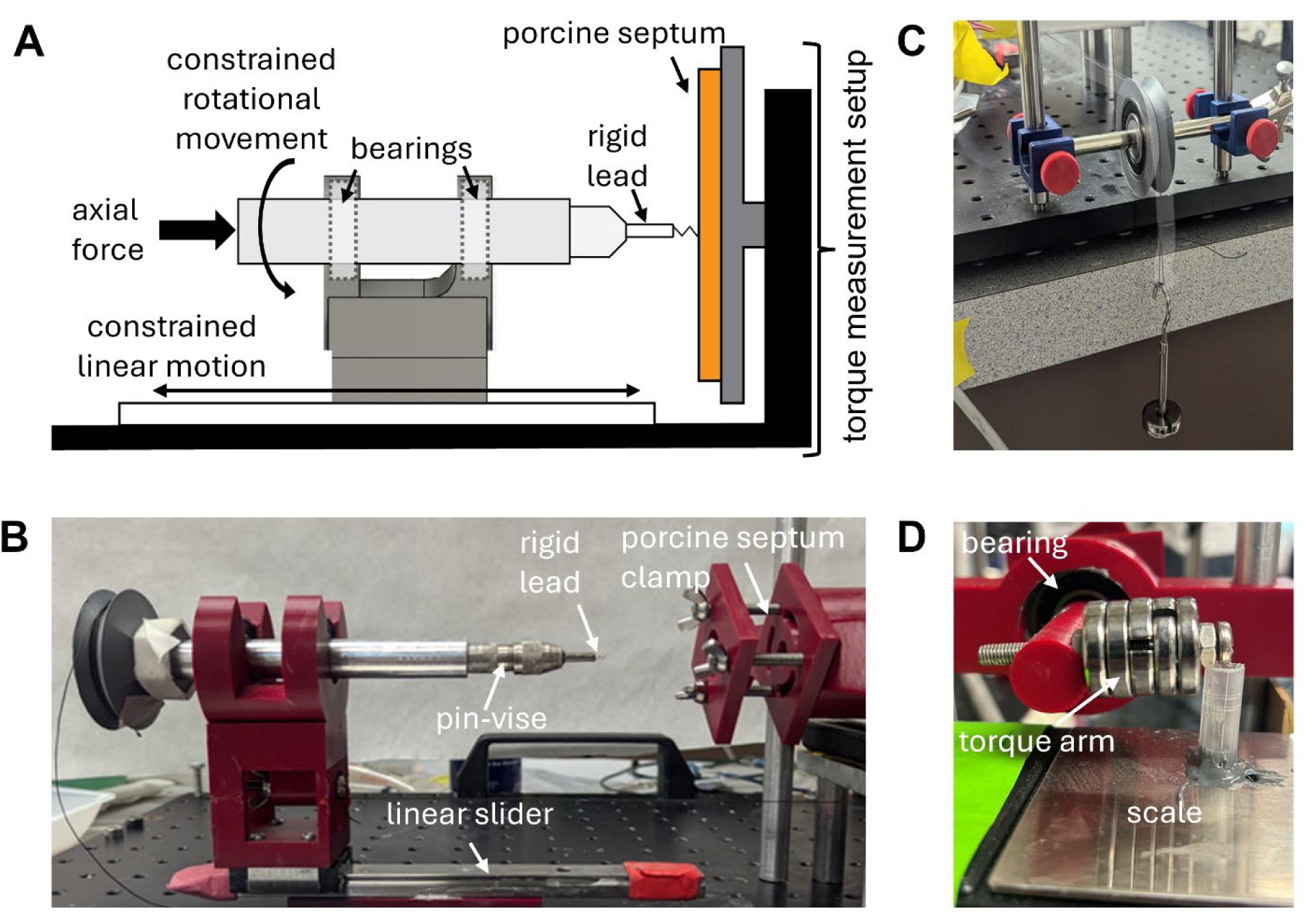
Idealized one-one lead rotation system used to assess lead insertion behavior and measure the muscle torque on a clamped porcine septal sample. **(A)** Schematic and **(B)** physical setup of the idealized rotation response system, which uses a linear slider to translate rotation to constrained linear motion. **(C)** Rigid-lead setup was interfaced with a pulley and weights to evaluate the impact of variable axial force on helix-muscle interaction. **(D)** Muscle torque measurement setup: the clamped septal sample is coupled to a scale via a bearing and torque arm preloaded with 50g.

An axial pre-load force was applied to the RLs to simulate the effect of differences in lead body design on helix-septal tissue interaction (**Figure 2A, 2C**). Two experienced clinical electrophysiologists (KY and SJ) provided benchtop estimates of axial force applied by a lumenless lead and a stylet-driven lead (Supplementary Figure A). Axial force to simulate lumenless leads (30g) and stylet-driven leads (60g) was applied to the connector holding RL on the linear slider using a pulley and weights. The RL linear slider assembly was gently pressed against the clamped septal sample prior to applying rotations. A total of 14 rotations were applied to the idealized rotation response system by manually pulling a string over period of approximately 50 seconds (Figure 2C). An average of 13 rotations has been reported during the final lead insertion attempt into the left bundle branch area^8^.

A custom torque measurement setup was developed to record the torque experienced by the muscle (in real-time) during each RL insertion attempt. The septal clamp was mechanically coupled to a digital scale (200g range; 0.01g resolution; Weigh Gram, CN) through a ball bearing (model 99502H-2RS; uxcell, CN) and a torque arm preloaded with 50g (Figure 2D). The load cell within the scale was interfaced with the cable of a DC bridge transducer amplifier (Supp Fig 2, ML110 BridgeAmp, AD Instruments, Colorado Springs, CO, USA). The amplified analog signal was recorded using LabChart software through a PowerLab 16/30 data acquisition system (AD Instruments, Colorado Springs, CO, USA).

This setup enabled torque measurements over a range of ±10Nmm. The accuracy of the measurement setup was verified before and after each RL experiment, at each axial force level, by applying known torques using weights (2 – 23g) suspended from the clamp via a string and pulley arrangement. A linear regression was fitted to the measured voltages corresponding to the applied torque, and the resulting calibration equation was used to convert voltage signals to torque values through this study (see Supplementary Figure B).

### 2.2 Development of rigid leads with variable helix designs

Helical coils with variable outer diameter (OD, 1mm, 2mm) and helix turns (2.25, 4.5) were fabricated by tightly wrapping stainless steel wires (0.012” dia., 304v spring-tempered SS, Wytech Industries, NJ) to the desired pitch (1mm, 2mm) over a shaft fixed in a bench vise. These helices were attached to hypo-tubes (OD: 0.12” and 0.072” SS) using steel-reinforced epoxy (JB Weld, Georgia). Finally, a single 15-20° facet was machined at the helix tip using a resin cutting disc on a Dremel tool at medium speed, consistent with the helix tip design of lead models used in clinical practice. The three helix design variables (OD, turns, and pitch) were evaluated at two levels each for a total of eight helix designs, at two axial forces (Figure 3A) in this experimental study.

**Figure 3.**
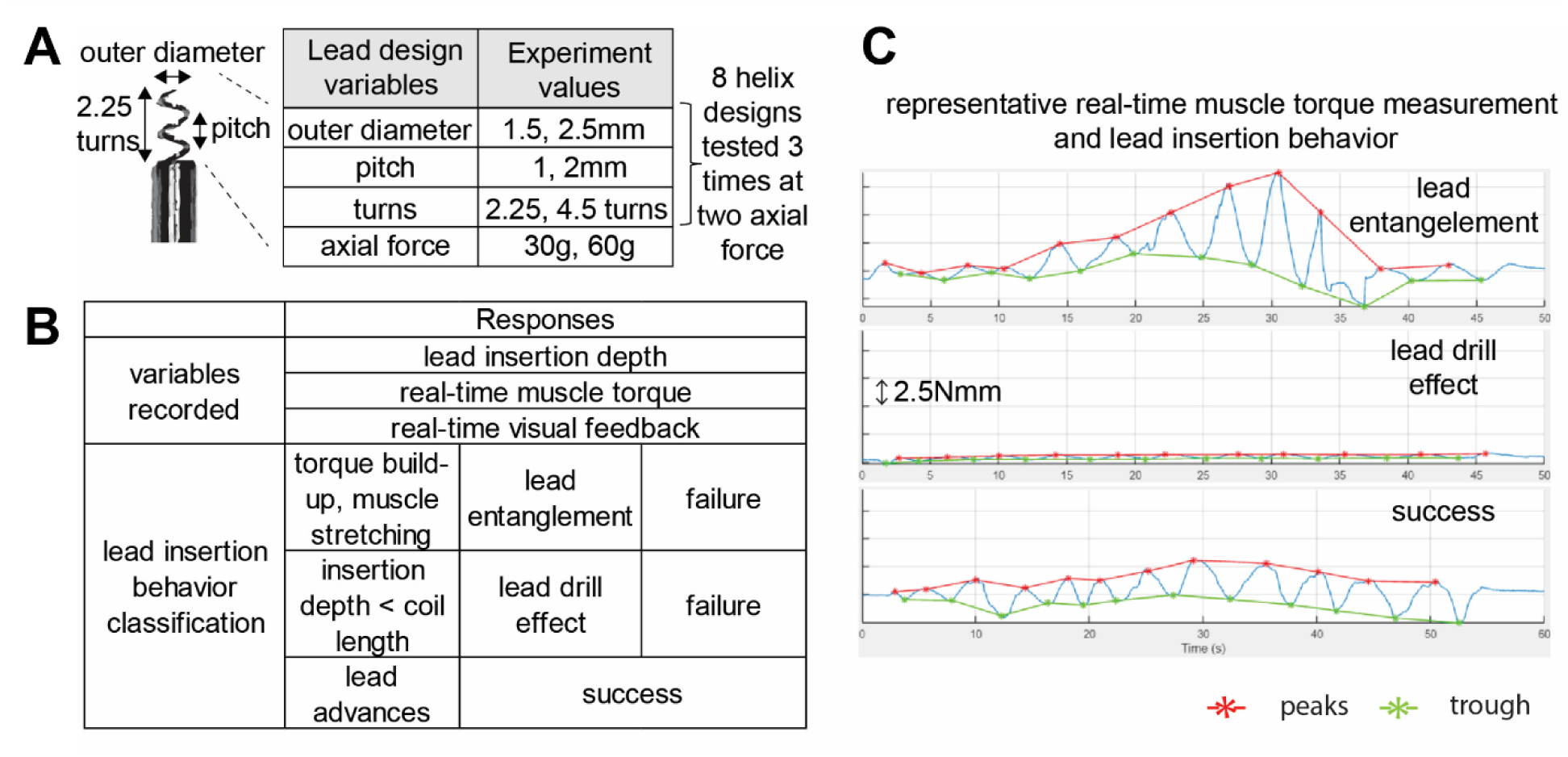
**(A)** Septal penetration behavior of rigid leads with eight helix designs (the outer diameter, pitch, and the number of helix turns were evaluated at two levels) were evaluated to two axial forces. **(B)** Three response variables were recorded: insertion depth (from displacement change after applying 14 rotations), real-time muscle torque of the clamped sample, and visual feedback (using camera). The three recordings were used to classify lead insertion behavior as successful or unsuccessful (entanglement or drill-effect). **(C)** Representative muscle torque traces used for classification of lead insertion behavior. Peaks and troughs (red/green asterisks) correspond to each rotation and were used to calculate the torque per turn (average of peak-peak values) and torque variability per turn (standard deviation of peak-peak values) for factorial analysis.

### 2.3 Experiment Plan

Freshly frozen porcine hearts (n=16, BCM Hearts, TX, USA) from 6-8 month old, 180-250lbs animals were defrosted in the refrigerator over 24 hours, and the IVS was excised. Anterior, posterior and apical tissue samples were prepared for RL insertion testing. Muscle torque, RL insertion depth, and video of the RL insertion attempt were recorded for analysis and classification of RL insertion behavior. Insertion was successful if the insertion depth exceeded the helix length and the muscle tissue was not entangled (Figure 3B). Video recording of muscle movement was used in conjunction with the real-time torque measurement to classify the lead behavior. Representative real-time muscle torque measurements for each lead behavior classification are shown in Figure 3C. Each helix design (n=8) was randomly inserted 3 times per axial force, and once per tissue sample. The experiment followed a blocked design by axial force, such that all insertions were first conducted under the 30g axial force and subsequently under the 60gaxial force, with the order of helix designs randomized within each block. The torque measurement setup accuracy was verified before and after experiments at each axial force, following the method outlined in a previous section (Experimental setup).

### 2.4 Statistical Analysis

A custom graphical user interface was developed in MATLAB (MathWorks, Natick, MA, USA) to pick the peaks and troughs corresponding to the RL rotations (‘*’ in red and green of Figure 3C, respectively). Notice the trend in the representative muscle torque traces for each classification of lead insertion behavior (Figure 3C). Lead advancement failure due to lead entanglement was characterized by larger amplitude muscle torque with RL rotations, whereas the “drill effect” was characterized by much smaller muscle torque amplitudes due to lack of lead-muscle engagement. Therefore, the mean of consecutive peak-peak amplitude of the muscle torque measurement (muscle torque per rotation), and the standard deviation of the consecutive peak-peak amplitude (muscle torque variability) was compared factorially.

Design factors’ effects on insertion success were analyzed using the chi-squared test, whereas the insertion depth was compared using the Mann-Whitney U Test. Design factors’ effect on muscle torque per rotation and muscle torque variability was also compared using the Mann-Whitney U-test. All aggregate values are reported as median (Q1-Q3). A P ≤ 0.05 was considered statistically significant.

## 3. Results

### 3.1 Outer Diameter and 30g Axial Force

At the 30g axial force setting, the larger OD RLs showed a trend toward improved insertion success (8/12 successful insertions) compared to the smaller OD RLs (5/12 successful insertions, P=0.22) (Figure 4A). Notably, the “drill effect” failure mode was observed only with the smaller OD and exclusively under this 30g axial force condition (2 out of 7 failed attempts). Larger OD RLs trended toward larger muscle torque per rotation as compared to smaller OD RLs (4.0Nmm, Q1-Q3: 3.77-4.87Nmm vs 3.53Nmm, Q1-Q3:3.24-3.74Nmm, P=0.17) (Figure 4B). The insertion depth of the successful septal lead penetration attempts was similar between the smaller and larger ODs at the 30g axial force (0.85cm, Q1-Q3: 0.80-1.02cm vs 0.99, Q1-Q3: 0.81-1.28, P=0.55) (Figure 4C).

**Figure 4.**
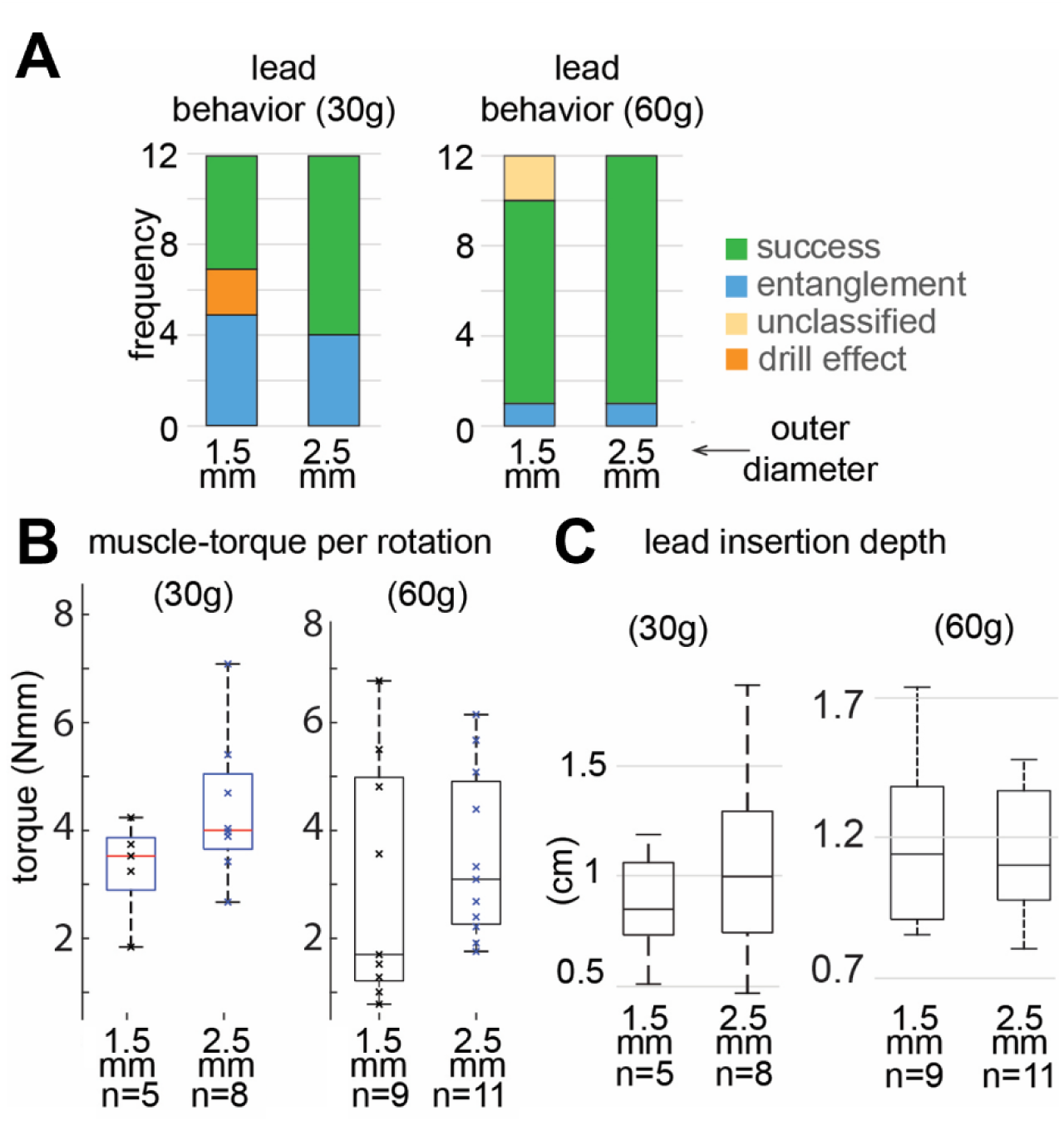
Impact of lead outer diameter (OD) on insertion behavior. **(A)** At 30g axial force, larger OD helices trended towards increased insertion success. The drill effect was observed only on smaller OD and only at the 30g axial force setting. **(B)** At 30g axial force, larger OD helices trended towards increased muscle torque per rotation. **(C)** OD did not seem to affect the insertion depth at the 30g or the 60g axial force setting.

### 3.2 Outer Diameter and 60g Axial Force

Increasing the axial force largely mitigated the differences between the smaller and larger OD RLs (9/10 vs 11/12 successes, P=0.89) (Figure 4A) observed at 30g. Muscle torque per rotation between the smaller and larger OD RL sizes showed no significant difference (P=0.29), with the smaller OD having a smaller median torque (1.64Nmm, Q1-Q3:1.11-4.84Nmm) than the larger OD (3.00Nmm, Q1-Q3: 2.35-4.69Nmm) (Figure 4B). The insertion depth of the successful septal lead penetration attempts was also similar between the smaller and larger ODs at the 60g axial force (1.14cm, Q1-Q3: 0.91-1.35cm vs 1.10, Q1-Q3: 0.99-1.33, P=1) (Figure 4C).

### 3.3 Pitch and 30g Axial Force

At the 30g axial force setting, the smaller pitch RLs trended towards improved insertion success compared to the larger pitch helices (8/12 vs 5/12, respectively, P=0.22) (Figure 5A). The “drill effect” failure mode was observed only with the larger pitch and only under the 30g axial force setting (2 out of 7 failed attempts). Smaller pitch RLs trended towards larger muscle torque per rotation (4.0Nmm, Q1-Q3: 3.85-4.53Nmm vs 3.42Nmm, Q1-Q3:3.24-3.53Nmm, P=0.21) (Figure 5B). The median insertion depth of the successful RL insertion attempts was lower for the smaller pitch RLs compared to the larger pitch RL at the 30g axial force setting (0.87cm, Q1-Q3: 0.57-1.05cm vs 1.19, Q1-Q3: 1.02-1.55, P=0.05) (Figure 5C).

**Figure 5.**
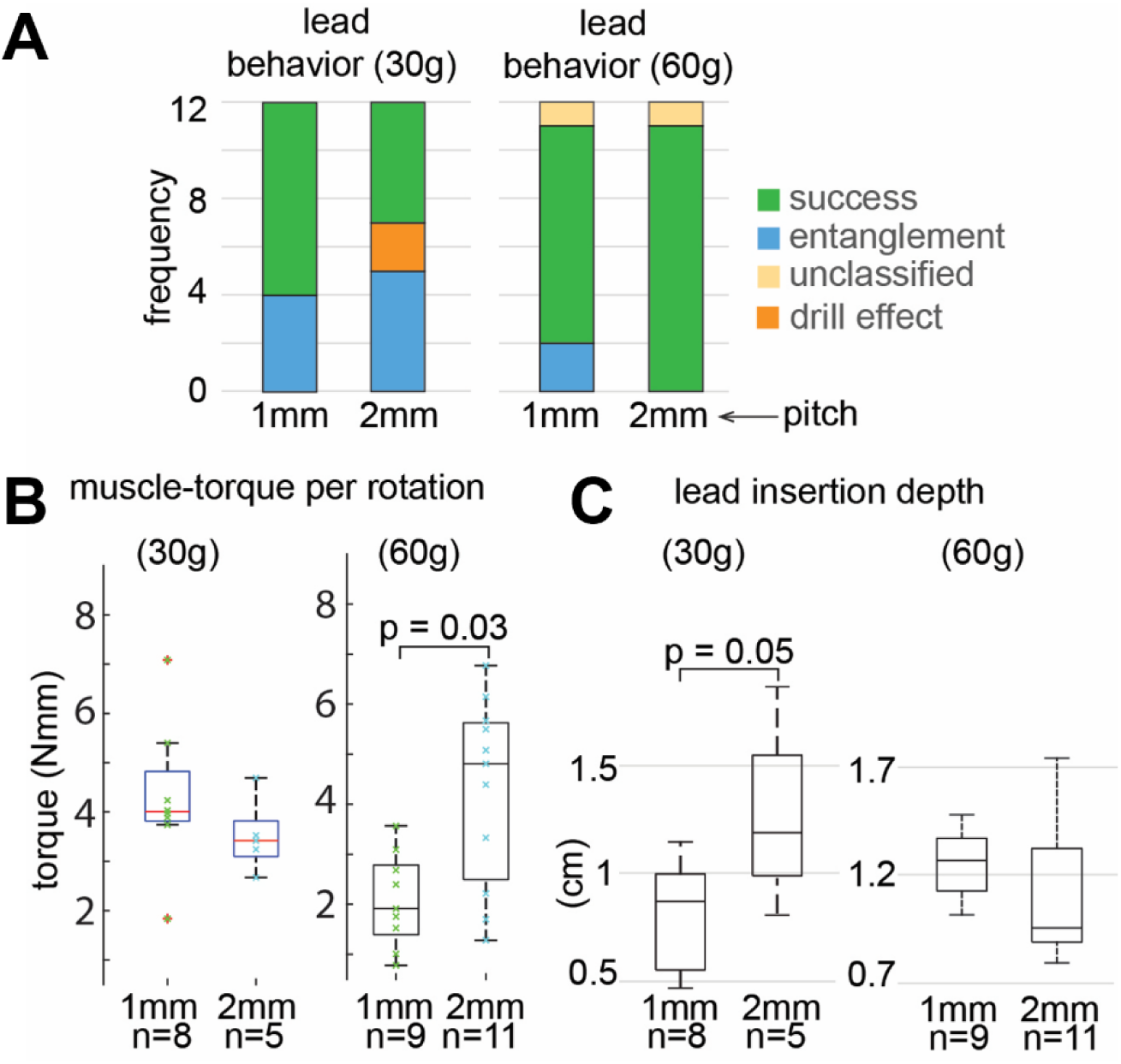
Impact of lead pitch on insertion behavior. **(A)** Leads with a smaller pitch trended towards increased insertion success. The drill effect was observed only on leads with a larger pitch and only at the 30g axial force setting. **(B)** Muscle torque per rotation showed conflicting results: at the 30g axial force, smaller pitch leads trended towards higher muscle torque per rotation. However, at the 60g axial force setting, larger pitch correlated to increase in muscle torque per rotation. **(C)** Leads with larger pitch achieved greater insertion depth at the 30g axial force setting. However, this difference was not significant at the 60g axial force.

### 3.4 Pitch and 60g Axial Force

Increasing the axial force largely mitigated the differences between the smaller and larger pitch RLs (9/11 vs 11/11, respectively, P=0.14) (Figure 5A) observed at 30g. Interestingly, for successful 60g axial force insertion attempts, smaller pitch RLs had a lower median muscle torque per rotation than the larger pitch RLs (1.96Nmm, Q1-Q3: 1.64-2.58Nmm vs 4.84Nmm, Q1-Q3:2.65-5.63Nmm, P=0.03) (Figure 5B). Furthermore, the median insertion depth of the successful RL insertion attempts trended larger for the smaller pitch helices compared to the larger pitch helices at the 60g axial force (1.27cm, Q1-Q3: 1.14-1.35cm vs 0.96, Q1-Q3: 0.9-1.25, P=0.08) (Figure 5C).

### 3.5 Helix turns and 30g Axial Force

At the 30g axial force setting, the RLs with fewer helix turns had a lower insertion success compared to RLs with more turns (4/12 vs 9/12, P=0.04) (Figure 6A, 6B). The “entanglement” mode of RL insertion failure occurred more often in RLs with fewer helix turns than in those with more turns (7/12 vs 2/12, P = 0.04). During successful insertions, no significant difference was noted for the muscle torque per rotation between the RLs with fewer helix turns vs more turns (5.05Nmm, Q1-Q3: 3.98-5.82Nmm vs 3.74Nmm, Q1-Q3:3.42-3.97Nmm, P=0.7) (Figure 6C). The median insertion depth of the successful RL insertion attempts trended lower for the fewer-turn helices compared to the helices with more turns at the 30g axial force (0.55cm, Q1-Q3: 0.50-0.80cm vs 1.02, Q1-Q3: 0.95-1.22, P=0.12) (Figure 6D).

**Figure 6.**
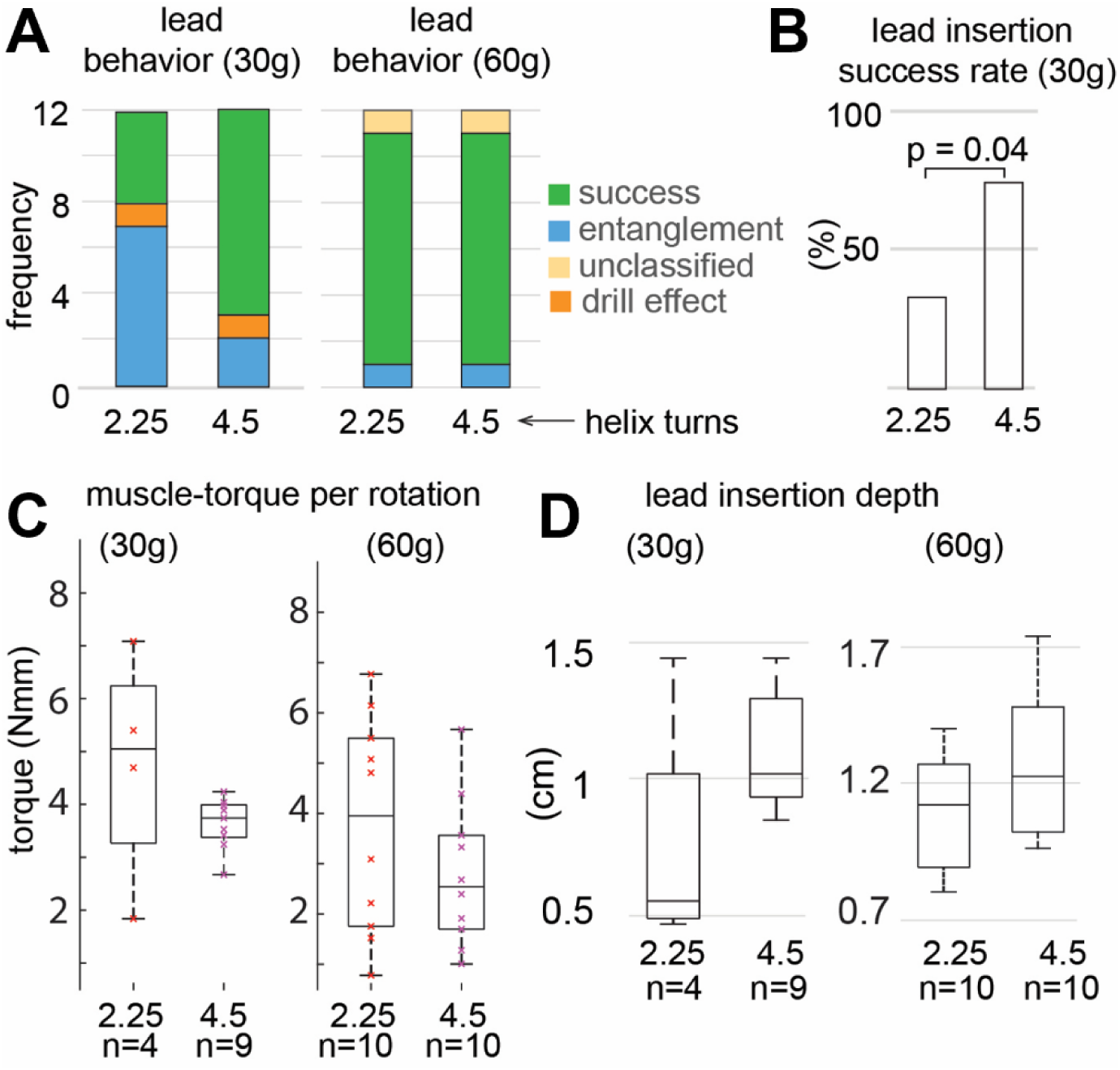
Impact of number of helix turns on lead insertion behavior. **(A)** and **(B)** At 30g axial force setting, leads with fewer helix turns failed to penetrate the septum more often due to entanglement, while leads with greater number of turns had a higher insertion success rate. The drill effect was only observed at this 30g axial force setting. **(C)** Muscle torque per rotation and (D) insertion depth were not significantly impacted by the number of helix turns at either the 30g or 60g axial force setting.

### 3.6 Helix Turns and 60g Axial Force

Increasing the axial force largely mitigated the differences between the RLs with fewer vs RLs with more helix turns (10/11 vs 10/11, respectively, P=1) observed at the lower setting (Figure 6A). No significant difference was noted between the fewer turn vs more turn RL muscle torque per rotation (3.97Nmm, Q1-Q3: 1.94-5.47Nmm vs 2.50Nmm, Q1-Q3:1.52-3.46Nmm, P=0.31) (Figure 6C). The median insertion depth of the successful septal lead penetration attempts trended lower for the fewer-turn helices compared to the helices with more turns at the 30g axial force (1.12cm, Q1-Q3: 0.89-1.26cm vs 1.23, Q1-Q3: 1.02-1.47, P=0.15) (Figure 6D).

### 3.7 Impact of Axial force

Higher (60g) axial force generally improved insertion success rate compared to insertion attempts at the 30g axial force (90.90% vs 54.17%, P = 0.006) (Figure 7A). Successful insertion attempts at 60g axial force showed trend towards an increased insertion depth compared to 30g axial force insertion depths (1.12cm Q1-Q3:0.96-1.37 vs 0.97cm Q1-Q3:0.80-1.14cm P=0.07) (Figure 7B).

**Figure 7.**
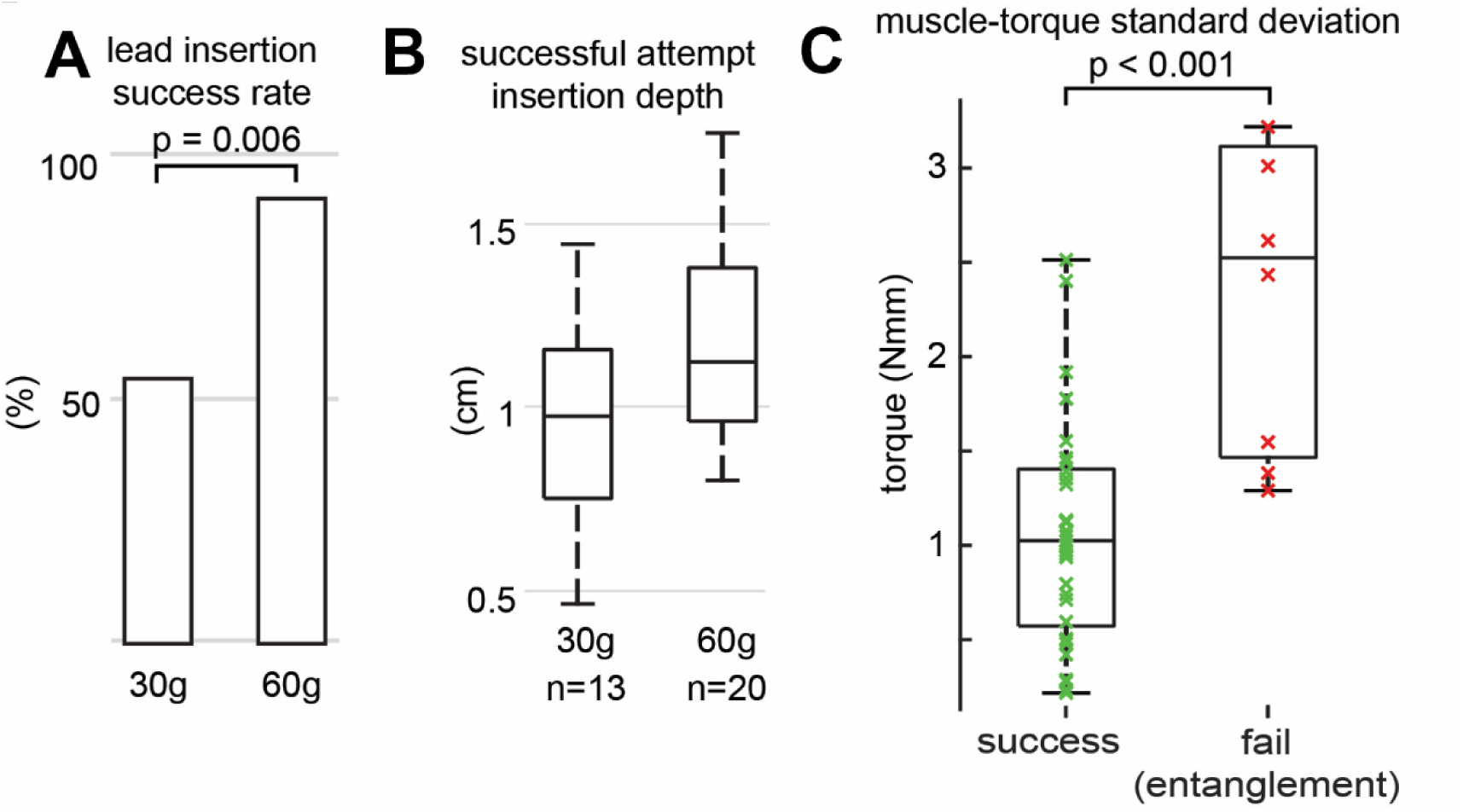
**(A)** 60g axial force resulted in more successful insertion attempts. **(B)** However, axial force did not significantly impact the lead insertion depth. **(C)** The muscle torque variability was significantly higher for the entangled insertion attempts compared to the successful insertion attempt cases.

The muscle torque variability for the successful cases was significantly smaller than the entangled mode of lead insertion failure (1.03Nmm Q1-Q3:0.6-1.39Nmm vs 2.53Nmm Q1-Q3: 1.51-3.06Nmm, P<0.001) (Figure 7C).

## 4. Discussion

The key findings of this study provide insights about the interplay among helix geometry, axial force, and lead insertion success. The observations highlight distinct mechanical behaviors at 30g versus 60g axial forces, which have implications for lead design and ease of use.

1. Number of helix turns: At the 30g axial force, RLs with a greater number of helix turns showed a high lead insertion success rate (Figure 6B). In contrast, RLs with fewer turns frequently exhibited the entangled mode insertion failure (Figure 6A).
2. Pitch and insertion depth: At the 30g axial force, RLs with smaller pitch tended to generate higher muscle torque per rotation and demonstrated a tendency to improve the overall success rate (Figure 5A). However, under the same conditions, the successfully inserted larger pitch RLs had achieved a deeper insertion depth. Interestingly, at the 60g axial force, RLs with larger pitch had larger muscle torque per rotation (Figure 5B).
3. Outer diameter: At the 30g axial force, larger outer diameter showed a trend towards increased muscle torque per rotation and increased success rate (Figure 4A, 4B).
4. Drill effect: The ‘drill effect’ mode of RL insertion failure was observed only at the 30g axial force condition and occurred in RLs with smaller OD (Figure 4A) and larger pitch (Figure 5A).
5. Muscle torque variability: The entangled mode of insertion failure was characterized by a significant amount of variability in the muscle torque over the duration of the insertion attempt (Figure 7C).

These observations may be interpreted with an intuitive mechanical framework.

Based on the torque relationship (Torque = Force x Radius), a helix with larger OD subjected to the same rotational force will generate more torque on the septal sample.

This theoretical relationship aligns with the observed trend of higher muscle torque per rotation and increase in insertion success rate for larger OD RLs.

Helix angle also plays a role. The angle between the septal surface and the helix coil is smaller in coils with a smaller pitch, allowing a greater portion of the torque to be transferred through the coils compared to coils with a larger pitch. Conversely, the angle between the axial force (perpendicular to the rotation direction) and the helix coil is smaller in coils with a larger pitch. Therefore, when axial force is applied, torque transfer is more efficient in coils with a larger pitch. This theoretical relationship is observed in the experimental data: at 30g axial force, smaller pitch RLs showed a trend towards higher muscle torque per rotation and greater insertion success. However, at the 60g axial force, the larger pitch helices had a higher muscle torque per rotation.

The RL insertion failure ‘drill effect’ occurs when the lead is stuck in a septal recess and does not engage with the myocardium. Only two instances of drill effect were observed – both were at the 30g axial force setting, involving RLs with a smaller OD, and a larger pitch. The 30g axial force likely reduced the helix-tissue contact, and the smaller OD helix is more prone to getting stuck within the septal recess. Additionally, the larger pitch likely reduced torque transfer, together explaining the observed behavior.

Longer helices have been proposed to more effectively pull the lead deeper into the septal tissue^12^. This trend was supported by the data; at the 30g axial force, RLs with a greater number of helix turns showed a higher insertion success rate. Under the same conditions, RLs with fewer turns exhibited a greater tendency toward the entangled mode of lead insertion failure.

Lumenless leads like the SelectSecure 3830 are thin, flexible, and generate lower axial force (Supplementary Figure A). As a result, they are more prone to entanglement, drill effect, and may fail to advance into thicker septa^7,10^. In contrast, stylet-driven leads provide higher axial force, and prior studies - consistent with the findings of this study - have reported shorter procedure times with these leads^13^. Therefore, this study offers important insights into helix coil configuration for future lead development.

## 5. Limitations

### 5.1 Ex Vivo Tissue Model and Preparation

This study was conducted on excised samples of the right ventricular septum that were obtained from freshly frozen porcine hearts. The samples were thawed in a refrigerator for over 24 hours. While this method of thawing has been shown to be effective for preserving the mechanical integrity of muscle tissue and yields properties closest to the freshly excised sample ^14^, it does not replicate the in vivo setting. The model lacks blood flow, and the dynamic environment of a beating heart. Despite these limitations, the relative comparison of leady body and helix variables within this ex vivo model remains relevant for establishing the mechanical principles for lead implantation.

### 5.2 Experimental Design and Measurement Constraints

This study exclusively evaluated the impact of lead body and helix variables on septal insertion. No direct feedback was recorded to identify if the lead had perforated the septal samples, which limits the interpretation of the final insertion depth and the torque measurements. Furthermore, the methods used in this study were insufficient to fully classify lead behavior related to the endocardial barrier effect. This was primarily because the insertion depth was quantified as the total translational displacement of the RL after applying rotations.

### 5.3 Procedural Differences and Data Exclusion

The application of rotations differed from the clinical practice. Clinically, 3-5 rapid rotations are recommended at a time. However, the 14 rotations applied in this study were applied over a period of 50 seconds. Given that all other mechanical factors were held constant across all the test groups, we assume the rotation speed did not significantly influence the outcomes of this experimental study.

Two RL insertion attempts at the 60g axial force setting, and of smaller OD, were excluded from the analysis. In these instances, the RL linear slider assembly accelerated toward the sample instead of gently pressing against it and preemptively punctured the septum. A trend of deeper insertion depth was observed at the 60g axial force setting (Figure 7A). There is a possibility that the lead dragged during rotation, although the factor cannot be confirmed or excluded with the methods used.

### 5.4 Scope Limitation

Finally, this study exclusively evaluated the impact of lead design factors to penetrate the septum. Our observations suggest a likely relationship between lead design factors and lead extractability. However, the evaluation of factors affecting lead extraction ability was outside the scope of this investigation.

## 6. Conclusions

Deep septal advancement of pacing leads often fails due to technical issues or due to scarred muscle. Research and development efforts are underway to address these unmet needs for left bundle branch area pacing^15,16^. In this work, we developed a rigid rotation-to-translation system to elucidate how helix geometry and axial force interact during lead insertion. Our findings show that 30g and 60g axial forces produce distinct insertion behaviors, each influenced by specific helix parameters. These insights will inform the design of future leads and support physicians in evaluating next-generation devices for clinical use.

## 7. Figures

## 8. Supplementary Figures

**Supplementary Figure A.**
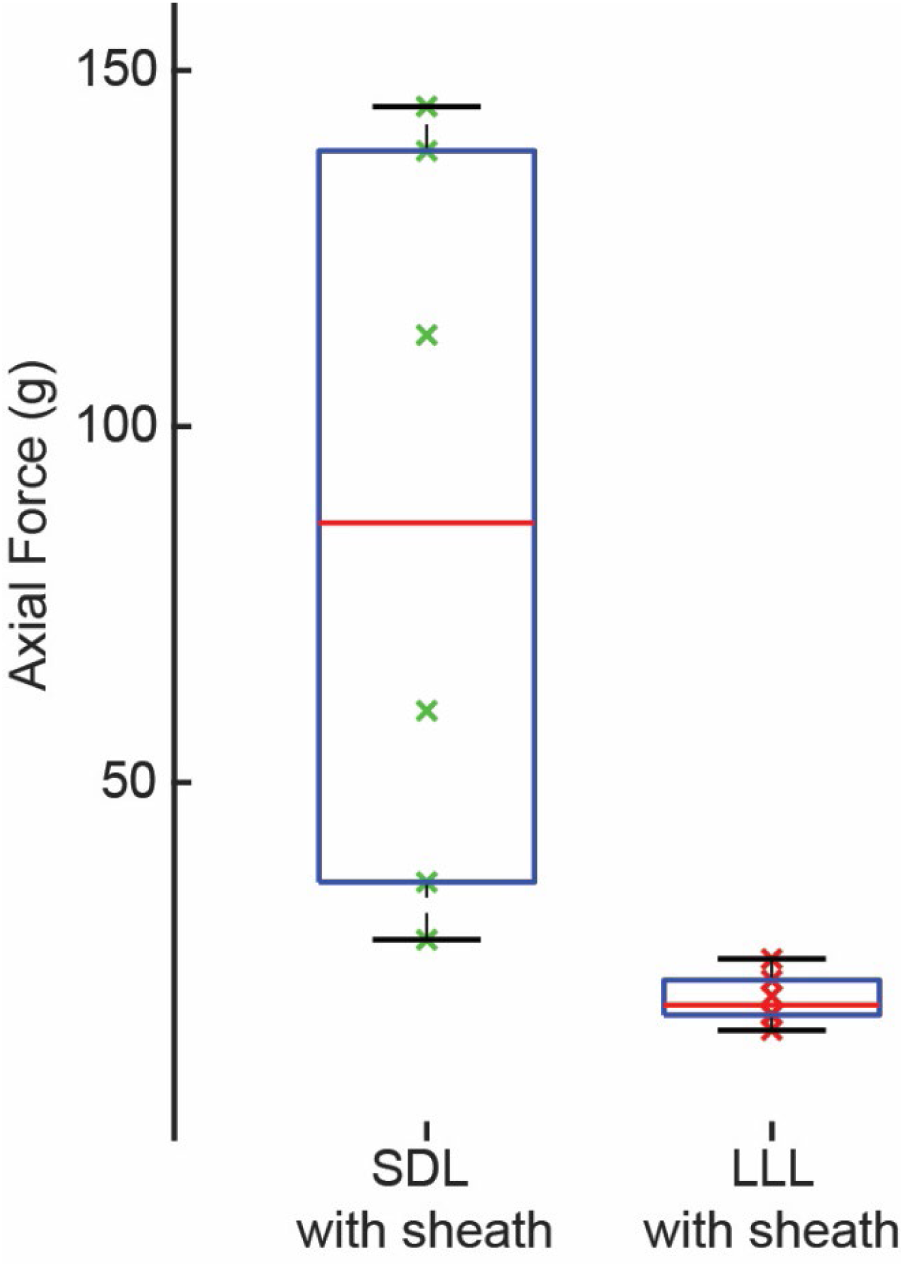
Axial force measurements using a stylet-driven lead (SDL, 5076 CapSureFix Novus MRITM, Medtronic, Inc) and a lumenless lead (LLL, 3830 SelectSecureTM, Medtronic, Inc). Two experienced electrophysiologists provided three repeated axial force measurements for the two lead types.

**Supplementary Figure B.**
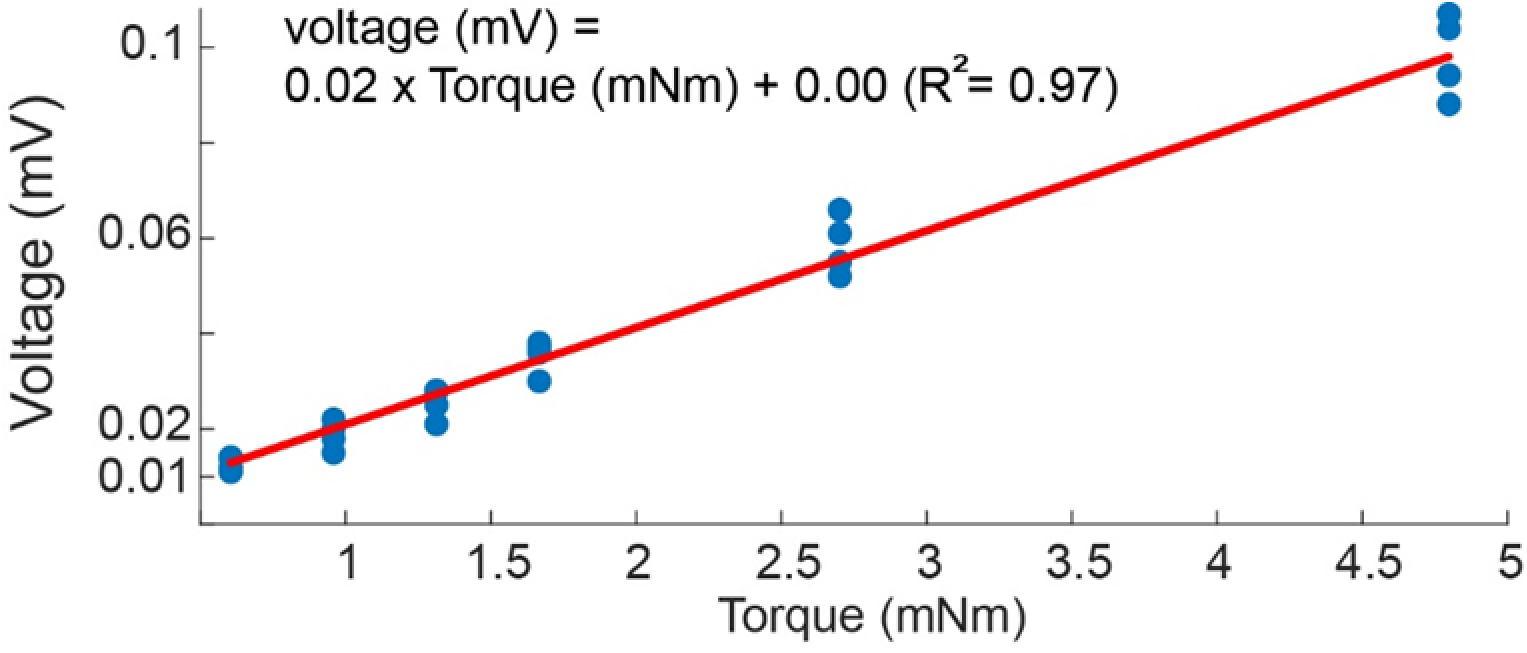
Linear regression for torque conversion. A linear regression was fitted to sensor voltages measured at known torques, using data collected before and after the experiment for each axial force setting. The resulting equation was used to convert all voltage signals to final torque values.

## Abbreviations

RL: Rigid Leads
OD: Outer diameter

## References

1. Chardack WM, Gage AA, Schimert G, Thomson NB, Sanford CE, Greatbatch W: Two years’ clinical experience with the implantable pacemaker for complete heart block. Dis Chest Elsevier, 1963; 43:225–239.

2. Mond HG, Helland JR, Stokes K, Bornzin GA, McVENES R: The electrode-tissue interface: The revolutionary role of steroid-elution. Pacing and Clinical Electrophysiology Wiley Online Library, 2014; 37:1232–1249.

3. Manolis AS, Simeonidou E, Sousani E, Chiladakis J: Alternate sites of permanent cardiac pacing: A randomized study of novel technology. Hellenic J Cardiol 2004; 45:147–151.

4. Glikson M, Burri H, Abdin A, et al.: European Society of Cardiology (ESC) clinical consensus statement on indications for conduction system pacing, with special contribution of the European Heart Rhythm Association of the ESC and endorsed by the Asia Pacific Heart Rhythm Society, the Canadian Heart Rhythm Society, the Heart Rhythm Society, and the Latin American Heart Rhythm Society. EP Europace [Internet] 2025; 27:euaf050. Available from: 10.1093/europace/euaf050

5. Burri H, Jastrzebski M, Cano Ó, et al.: EHRA clinical consensus statement on conduction system pacing implantation: endorsed by the Asia Pacific Heart Rhythm Society (APHRS), Canadian Heart Rhythm Society (CHRS), and Latin American Heart Rhythm Society (LAHRS). Europace Oxford University Press, 2023; 25:1208–1236.

6. Sritharan A, Kozhuharov N, Masson N, Bakelants E, Valiton V, Burri H: Procedural outcome and follow-up of stylet-driven leads compared with lumenless leads for left bundle branch area pacing. Europace Oxford University Press, 2023; 25.

7. Jastrzębski M, Moskal P, Hołda MK, et al.: Deep septal deployment of a thin, lumenless pacing lead: a translational cadaver simulation study. EP Europace Oxford University Press, 2020; 22:156–161.

8. Zou J, Chen K, Liu X, et al.: Clinical use conditions of lead deployment and simulated lead fracture rate in left bundle branch area pacing. J Cardiovasc Electrophysiol Wiley Online Library, 2023; 34:718–725.

9. Chapman D, Morgan F, Tiver KD, et al.: Assessing Torque Transfer in Conduction System Pacing: Development and Evaluation of an Ex Vivo Model. Clinical Electrophysiology American College of Cardiology Foundation Washington DC, 2024; 10:306–315.

10. Valappil SP, Chapman D, Muenzinger C, et al.: Left bundle area pacing in hypertrophied hearts: An ex vivo ovine model to study deployment of pacing leads in thick septum. Heart Rhythm [Internet] Elsevier, 2025; 22:2055–2064. Available from: https://linkinghub.elsevier.com/retrieve/pii/S1547527125023914

11. Grosfeld MJW, Vos DHS, De Boer TJM, Bos HJ: Testing a new mechanism for left interventricular septal pacing: The transseptal route: A feasibility and safety study. Europace Oxford University Press, 2002; 4:439–444.

12. Mafi-Rad M, Luermans JGLM, Blaauw Y, et al.: Feasibility and acute hemodynamic effect of left ventricular septal pacing by transvenous approach through the interventricular septum. Circ Arrhythm Electrophysiol Am Heart Assoc, 2016; 9:e003344.

13. Sripusanapan A, Wareesawetsuwan N, Deepan N, et al.: Comparing Efficacy and Complications Between Stylet-Driven Leads and Lumenless Leads in Left Bundle Branch Area Pacing. Pacing and Clinical Electrophysiology Wiley Online Library, 2025; .

14. Park MH, Kim M: Effects of thawing conditions on the physicochemical and microbiological quality of thawed beef. Prev Nutr Food Sci 2024; 29:80.

15. Shah AR, Yazaki K, Seamons B, et al.: Multiple electrode leads facilitate left bundle branch area pacing: A concept evaluation study. Heart Rhythm O2 [Internet] Elsevier, 2025; 6:1005–1010. Available from: 10.1016/j.hroo.2025.04.012

16. Reddy VY, Nair DG, Doshi SK, et al.: First-in-human study of a leadless pacemaker system for left bundle branch area pacing. Heart Rhythm Elsevier, 2025; .

